# Temperature response of Rubisco kinetics in *Arabidopsis thaliana*: thermal breakpoints and implications for reaction mechanisms

**DOI:** 10.1101/320879

**Authors:** Ryan A. Boyd, Amanda P. Cavanagh, David S. Kubien, Asaph B. Cousins

## Abstract

Optimization of Rubisco kinetics could improve photosynthetic efficiency, ultimatly resulting in increased crop yield. However, imprecise knowledge of the reaction mechanism and the individual rate constants limit our ability to optimize the enzyme. Membrane inlet mass spectrometery (MIMS) may offer benefits over traditional methods for determining individual rate constants of the Rubisco reaction mechanism, as it can directly monitor concentration changes in CO_2_, O_2_, and their isotopologs during assays. However, a direct comparsion of MIMS to the traditional Radiolabel method of determining Rubisco kinetic parameters has not been made. Here, the temperature responses of Rubisco kinetic parameters from *Arabidopsis thaliana* were measured using the Radiolabel and MIMS methods. The two methods provided comparable parameters above 25 °C, but temperature responses deviated at low temperature as MIMS derived catalytic rates of carboxylation, oxygenation, and CO_2_/O_2_ specificity showed thermal breakpoints. Here we discuss the variability and uncertainty surrounding breakpoints in the Rubisco temperature response and relavance of individual rate constants of the reaction mechanisms to potential breakpoints.

## INTRODUCTION

The enzyme Ribulose-1,5-bisphosphate carboxylase/oxygenase (Rubisco) catalyzes the reaction of CO_2_ or O_2_ with Ribulose-1,5-bisphosphate (RuBP) initiating the photosynthetic carbon reduction cycle or photorespiratory cycle, respectively (Bowes *et al*., 1971; Andrews *et al*., 1973). Kinetic studies on Rubisco typically report the Michaelis-Menten constants for carboxylation (*K*_C_) and oxygenation (*K*_O_), the catalytic rate of carboxylation (*k*_catCO2_) and oxygenation (*k*_catO2_), and the specificity of the enzyme for CO_2_ over O_2_ (*S*_C/O_) as these parameters are used for modeling leaf gas exchange (von Caemmerer, 2000). Each of the above Michaelis-Menten parameters is a combination of elementary rate constants that describe the reaction mechanism; however, the rate constants are less well studied (Tcherkez, 2013; Tcherkez, 2016). Optimization of Rubisco kinetics for enhanced CO_2_ reduction has been proposed (Spreitzer and Salvucci, 2002), but this effort is limited by our current understanding of the reaction mechanism (Tcherkez *et al*., 2006; Tcherkez, 2013).

The carboxylation and oxygenation reaction mechanisms can be separated into elementary rate constant as originally proposed by Farquhar (1979), reviewed by Tcherkez (2013) and reproduced in Figure 1. Since the initial description of the reaction mechanism (Hurwitz *et al*. 1956) there has been slow progress in defining rate constants due to experimental difficulties in isolating their individual effects. However, the use of membrane inlet mass spectrometry (MIMS) to study Rubisco kinetics may hold promise. The traditional Radiolabel method used in most Rubisco publications relies on ^14^C assays to determine *k*_catCO2_, *K*_C_, *K*_O_, a separate ^3^H assay to determine *S*_C/O_, leaving *k*_catO2_ to be calculated. Alternatively, the MIMS assay simultaneously measures changing concentrations of CO_2_ and O_2_ and can therefore determine all kinetic parameters with a single assay (Cousins *et al*., 2010; Boyd *et al*., 2015). An advantage of the MIMS method is that in addition to the abundant isotopologues of CO_2_ (^12^CO_2_) and O_2_ (^16^O_2_) the system can monitor less abundant stable isotopologues such as ^13^CO_2_ and ^16^O^18^O. Measurements of primary kinetic isotope effects have been useful in defining enzyme reaction mechanisms (O’Leary *et al*., 1992); therefore, the MIMS system may provide new information regarding the individual rate constants. At 25 °C the MIMS method has been used for determining both Rubisco carbon fractionation (McNevin *et al*., 2006; McNevin *et al*., 2007; Tcherkez *et al*., 2013), and Michaelis-Menten constants of the carboxylation (*v*c) and oxygenation (*v*o) reactions (Cousins *et al*., 2010). Additionally, it was used to determine the temperature dependencies of the Rubisco kinetic parameters in the C_4_ species *Setaria viridis* (Boyd *et al*., 2015). However, previous work using the Radiolabel method suggest lower *E*_a_ values for *V*cm_a_x in C_4_ species than that measured by Boyd *et al*. (2015; Sharwood *et al*., 2016; Sage, 2002; Kubien *et al*., 2003; Perdomo *et al*., 2015), suggesting comparisons between the MIMS *E*_a_ values and the traditional Radiolabel method are needed.

**Figure 1.**
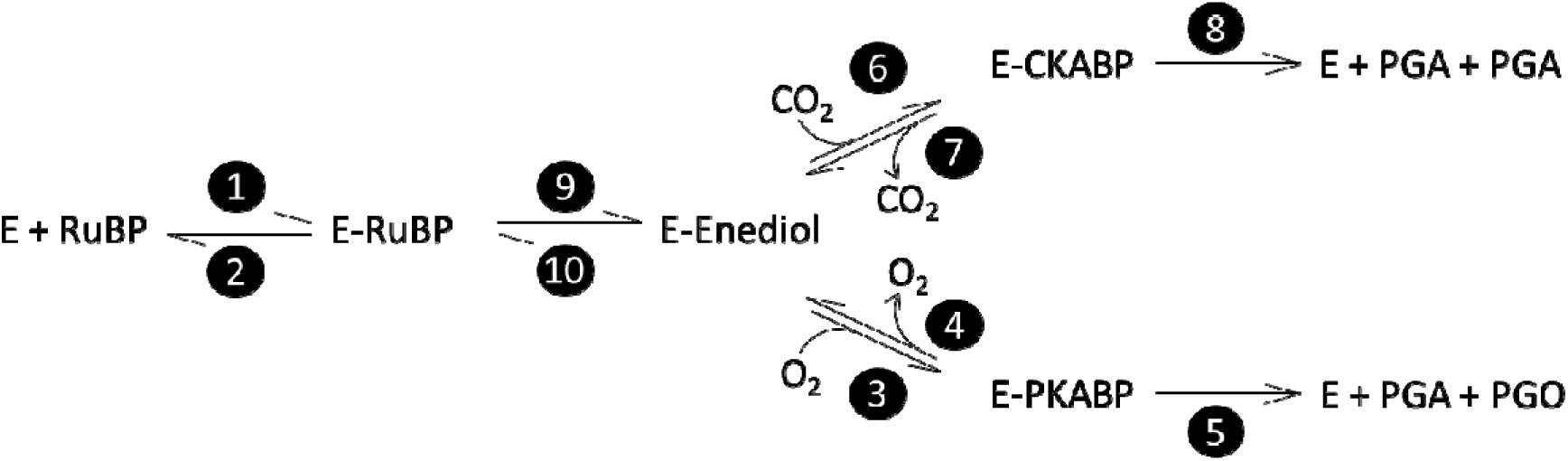
Elementary reactions of Rubisco-catalyzed carboxylation and oxygenation reproduced from Farquhar (1979). Each reaction, forward and reverse, is numbered in a filled circle following the numbering from Farquhar (1979). Steps 8 and 5 are written as irreversible reactions. Step 8 includes cleavage, hydration, and reprotonation as a single step. Step 5 includes cleavage and hydration as a single step. Each step is associated with a rate constant (*k*) and energy of activation (Δ*G*^‡^) following the same numbering as shown in filled circles. Abbreviations as follows: E, free activated enzyme; RuBP, D-ribulose-1,5-bisphosphate; E-RuBP; enzyme bound RuBP; E-Enediol, enzyme bound 2,3-enediolate form of RuBP; CO_2_, carbon dioxide; E-CKABP, enzyme bound carboxyketone intermediate; PGA, 3-phospho-D-glycerate; O_2_, oxygen; E-PKABP, peroxo intermediate; PGO, 2-phospho-glycolate.

Here we measured the temperature response of Rubisco kinetic parameters from *Arabidopsis thaliana* using two methods. First, we used the traditional method involving the use of radiolabeled substrate and analysis of labeled products following the reaction in known concentrations of CO_2_ and O_2_ (Jordan and Ogren, 1981), which we referred to as the Radiolabel method. Secondly, we used the MIMS method following the simultaneous consumption of CO_2_ and O_2_ over time, giving a direct measure of *v*c, *v*o, CO_2_, and O_2_ leading to simultaneous determination of *k*_catCO2_, *k*_catO2_, *K*_C_, *K*_O_, and *S*_C/O_ (Cousins *et al*., 2010; Boyd *et al*., 2015). Additionally, for the Radiolabel method we compared curve fitting CO_2_ responses to determine *K*_C_ and *k*_catCO2_ simultaniously in an O_2_ free buffer, and *k*_catCO2_ determined at a single bicarbonate concentration at all temperatures in open air. The later is a common practice for determining *k*_catCO2_ temperature responses (Tieszen and Sigurdson, 1973; Sage *et al*., 1995; Crafts-Brander and Salvucci, 2000; Pittermann and Sage, 2000; Sage, 2002; Kubien *et al*., 2003; Perdomo *et al*., 2015).

Recently, the existence of breakpoints in the *k*_catCO2_ temperature response was highlighted as a source of variability in the Rubisco temperature response literature (Sharwood *et al*., 2016). Initial observations of breakpoints in *V*cm_a_x temperature responses were determined to be a methodological artifact due to the use of a single bicarbonate concentration at all temperatures and were corrected by varying bicarbonate concentration with temperature (Björkman and Pearcy, 1970). However, breakpoints were later observed for *k*_catCO2_, *k*_catO2_, and *K*_C_ at 15 °C using a curve fitting method (Badger and Collatz, 1977). It was suggested that these breakpoints could be due to changes in rate limiting steps of the reaction mechanism caused by changes in enzyme conformation (Badger and Collatz, 1977). An additional breakpoint was reported in the *k*_catCO2_ of *Oryza sativa* at 22 °C (Sage, 2002) and Kubien *et al*. (2003) observed different temperature responses when *k*_catCO2_ was measured from 0 to 12 °C compared to 18 to 42 °C in *Flaveria bidentis*. Most recently, Sharwood *et al*. (2016) observed breakpoints in *k*_catCO2_ at 25 °C for Panicoid grasses when using a curve fitting method. Inconsistencies are evident between studies, and it is unclear if breakpoints are universal to all temperature response studies of plant Rubisco. Here, we discuss the possible causes of breakpoints, focusing on the three previously proposed causes of breakpoints: erroneous bicarbonate concentrations, changes in rate limiting step of the reaction mechanism, and deactivation of Rubisco at low temperature, using the Radiolabel and MIMS data sets reported here.

## MATERIALS AND METHODS

### Plant Growth

Plants for the Radiolabel method were grown and assayed at the University of New Brunswick, Fredericton, Canada. *Arabidopsis thaliana* (Col-0) seeds were stratified for 3 days at 4 °C on Promix (Plant Products, Brampton, Canada), transferred to a growth chamber (E-15, Conviron, Winnipeg, Manitoba, Canada) and grown under photoperiod conditions of 10 h light/14 h dark, 20/18 °C, and a photosynthetic photon flux density (PPFD) of 300 μmol m^−2^ s^−1^. Plants were watered with modified Hoagland’s solution as needed.

Plants for MIMS were grown and assayed at Washington State University, Pullman, Washington, U.S.A. Seeds of *A. thaliana*, ecotype Col-0, were placed in 2 L pots containing commercial soil (LC1 Sunshine Mix, Sun Gro Horticulture, Vancouver, Canada) and grown in an environmental growth chamber (Biochambers GC-16, Winnipeg, Manitoba, Canada) at a PPFD of 300 µmol m^−2^ s^−1^ at plant height, relative humidity was not controlled, air/night temperature of 23/18 °C, with a 14 hour photoperiod and 10 hours of dark. Plants were fertilized weekly (Peters 20-20-20, Allentown, PA, USA) and watered as needed.

### Sampling for Radiolabel Analysis

Leaf punches were obtained at mid-day, flash frozen in liquid nitrogen and stored at −80 °C until extraction. Leaf tissue was ground (1.1 cm^2^ disks, ca. 20 mg) in a Tenbroeck glass tissue homogenizer containing 3 mL of ice-cold extraction buffer (100 mM HEPES pH 7.6, 2 mM Na-EDTA, 5 mM MgCl_2_, 5 mM dithiothreitol [DTT], 10 mg ml^−1^ polyvinyl polypyrolidone, 2% (vol/vol) Tween-80, 2 mM NaH_2_PO_4_, 12 mM amino-*n*-capronic acid, and 2 mM benzamidine) and 50 μL Protease Inhibitor Cocktail (Sigma). This leaf homogenate was centrifuged at 16000× g at 4 °C for 60 seconds. The resulting supernatant was then desalted and concentrated as described by Cousins *et al*. (2010), and aliquots were incubated with 20 mM MgCl_2_ and 10 mM NaHCO_3_ at 30 °C for 20 min to fully carbamylate Rubisco. Rubisco content was quantified using [^14^C]carboxy-arabinitol bisphosphate (^14^CABP)-binding assay (Ruuska *et al*., 1998, Kubien *et al*., 2011).

### Sampling for MIMS Analysis

The youngest fully expanded leaves of plants 30 to 40 days after planting were sampled for Rubisco extraction. The mid vein was removed and approximately 2 g of leaf tissue was ground in 2 mL of ice-cold extraction buffer (100 mM HEPES pH 7.8, 10 mM DTT, 25 mM MgCl_2_, 1 mM EDTA, 10 mM NaHCO_3_, 1% (g/mL) PVPP, 0.5% (v/v) 100x Triton) with a mortar and pestle on ice. 67 µL of protease inhibitor cocktail (P9599, Sigma-Aldrich, St. Louis, Missouri) to 2 g of fresh leaf tissue was added to the extraction buffer before grinding. The homogenized extract was centrifuged at 4 °C, for 10 min, at 17000× g. The supernatant was collected and desalted using an Econo Pac 10DG column (Bio-Rad), filtered through a Millex GP 33-mm syringe-driven filter unit (Millipore), and then centrifuged using Amicon Ultra Ultracel 30K centrifugal filters (Millipore) at 4 °C for 1 hour at 3000× g. The layer maintained above the filter unit was collected, brought to 20% (v/v) glycerol, flash frozen in liquid nitrogen, and stored at −80 °C until measured. Rubisco content was determined as described above.

### Radiolabel Measurement of Rubisco Kinetic Parameters

The maximum carboxylation rate of fully activated Rubisco (*V*cm_a_x) was measured following the methods of Kubien *et al*. (2011) from 0 to 35 °C, by the incorporation of ^14^C into acid-stable products. This method is later referred to as the ‘single point’ method. Assays were initiated by the addition of 50 μL of activated extract (as described above) to 250 μL assay buffer (100 mM Bicine-NaOH (pH 8.2), 1 mM Na-EDTA, 20 mM MgCl_2_, 5 mM DTT, 400 μM RuBP, and 11 mM NaH^14^CO_3_ [~700 Bq nmol^−1^]), and stopped after 30 to 60 seconds by adding 250 μL of 1 M formic acid. Samples were dried at 90 °C, and ^14^C acid stable products were counted using a scintillation counter (LS-6500, Beckman-Coulter). Michaelis-Menten parameters for CO_2_ (*K*C), and apparent *K*_C_ at 21% O_2_ (*K*_C_ (21% O_2_)) were determined by assaying Rubisco activity in 7 mL septum-sealed, N2-sparged vials over a range of seven NaH^14^CO_3_ concentrations (Paul *et al*., 1991; Kubien *et al*. 2008. This analysis, referred to as the ‘curve fitting’ method, gave a separate temperature response of *k*_catCO2_ from the single point method described above. Rubisco *S*_C/O_ was determined following the method described by Kane *et al*. (1994). Additional details for these assays are presented in the supplemental files.

### MIMS Measurement of Rubisco Kinetic Parameters

Rubisco assays were conducted in a 600 µL temperature controlled cuvette linked to an isotope ratio mass spectrometer (Thermo-Fischer Delta V) and calibrated as previously described (Cousins *et al*., 2010; Boyd *et al*., 2015). Samples were measured similar to Boyd *et al*. (2015); four oxygen concentrations ranging from 40 to 1600 μM, and five CO_2_ concentrations ranging from 10 to 200 μM at each oxygen level were made. Measurements were made in 5 °C intervals from 10 °C to 40 °C, and the same three replicates were measured at each temperature. The assay buffer contained 200 mM HEPES (pH 7.7 at each measurement temperature), 20 mM MgCl_2_, 0.1 mM α-hydroxypyridinemethanesulfonic acid (α-HPMS), 8 mg mL^−1^ CA (Sigma), and 0.6 mM RuBP. 10 µL of extract was added per measurement. Rubisco was activated by leaving the extract at room temperature for 10 minutes prior to returning to ice before measurement. Additional details for these measurements are presented in the supplemental files.

### Modeling Temperature Responses

The temperature responses of the kinetic parameters were calculated for the equation

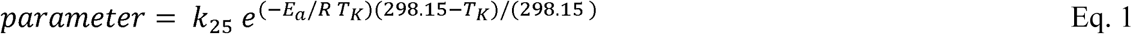

where *k*_25_ is the value of the parameter at 25 °C, *E*_a_ is the Arrhenius activation energy (kJ mol^−1^), *R* is the molar gas constant (0.008314 kJ mol^−1^ K^−1^), *T*_K_ is the temperature in Kelvin, and the term (298.15-*T*_K_)/298.15 is the scaling factor so that *k*_25_ may be used as the pre-exponential term. The *E*_a_ and *k*_25_ values for each Rubisco parameter were calculated by a linear regression of the natural log of the data plotted against (*T*_K_-298.15)/(*T*_K_), such that the y-intercept was equal to natural log of *k*_25_ and the slope was equal to *E*_a_/(298.15 *R*). For comparison the non-transformed temperature response are presented in Supp. Fig. 3. Three replicates of *E*_a_ and *k*_25_ were determined for each parameter, with the exception of Radiolabel *S*_C/O_ where the replication was four. For all MIMS and Radiolabel comparisons, other than *k*_catCO2_, only the curve fitting methods are compared. For simplicity we exclude the Radiolabel single point when comparing ratios of kinetic parameters to MIMS. Differences in the *k*_25_ and *E*_a_ values were determined by ANOVA, followed by pair-wise comparison (Tukey HSD) with a significance cutoff of p < 0.05 in Statistix 9 (Analytical Software, Tallahassee, USA).

Arrhenius plots for all kinetic parameters were examined for thermal breaks using the package ‘segmented’ in R, which first tests for differences between slopes using the Davies test (Davies 1987), and then estimates the breakpoints in linear models using maximum likelihood (Muggeo 2003; Muggeo 2008; R Core Team, 2013, http://www.R-project.org/). When breakpoints in the Arrhenius temperature response plots were statistically valid, the *E*_a_ values above and below the break points were compared to other *E*_a_ values as described above, the *k*_25_ value was held constant when fitting for two *E*_a_ values above and below the breakpoint.

### Equations for Reaction Mechanisms

Figure 1 depicts the currently hypothesized reaction mechanism of Rubisco as originally described by Farquhar (1979). The kinetic parameters *k*_catCO2_, *k*_catO2_, *K*_C_, *K*_O_, and *S*_C/O_ can be described by the individual first order rate constants (*k*) seen in Figure 1 as follows:

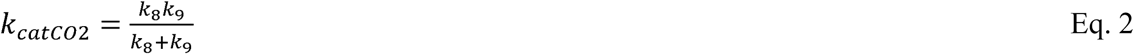

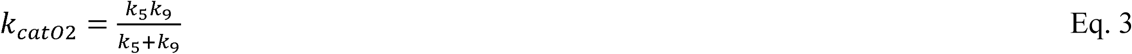

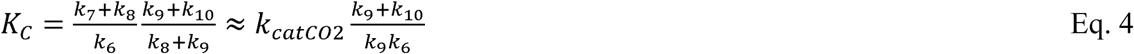

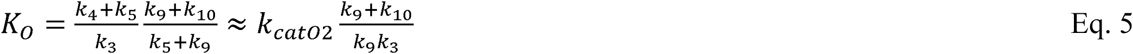

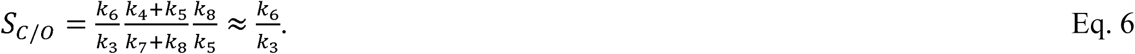

where the subscript indicates the transition state as numbered in Figure 1 by the black circles. The approximations in Equations 4, 5, and 6 are made by assuming the rates of decarboxylation (*k*_7_) and deoxygenation (*k*_4_) are negligible.

These first order rate constants can be related to temperature using transition state theory and the Eyring equation

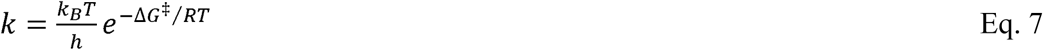

where *k*_B_ is the Boltzmann constant (1.3807·10^−23^ J K^−1^), *h* is the Planck constant (6.6261·10^−34^ J s), Δ*G*^‡^ (J mol^−1^) is the standard free energy difference between the transition state and the substrate (or intermediate). Note the proportionality constant *κ*, describing the proportion of vibrations that lead to product formation, has been assumed equal to one and left out of the equation. The Δ*G*^‡^ has components of entropy (Δ*S*^‡^) and enthalpy (Δ*H*^‡^) as defined by

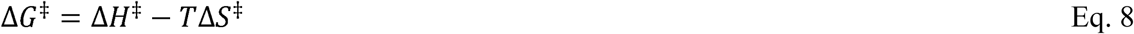

where the double dagger symbol (^‡^) denotes the transition state.

### Modeling *k* and Δ*G*^‡^

The proposed Rubisco reaction mechanism (Fig. 1) suggests *k*_catCO2_, *k*_catO2_, *K*_C_, *K*_O_, and *S*_C/O_ are described by complex terms made up of two or more elementrary reaction rates (Farquhar, 1979; Equations 2 through 6). The rate of an elementary reaction (*k*) is related to the energy barrier for the transition state of the reaction, often referred to as the activation energy (Δ*G*^‡^). The relationship between *k*, Δ*G*^‡^, and temperature is described by the Eyring equation (Equation 7), where Δ*G*^‡^ has enthalpic (Δ*H*^‡^) and entropic (Δ*S*^‡^) components (Equation 8). From Equation 8, a plot of Δ*G*^‡^ with temperature has a slope of Δ*S*^‡^ and a y-intercept of Δ*H*^‡^. For the discussion of Rubisco kinetics all numbering of *k*, Δ*G*^‡^, Δ*H*^‡^, Δ*S*^‡^ refer to reaction steps initially described by Farquhar (1979) and reproduced in Figure 1. The Eyring equation has been previously used to calculate Δ*G*^‡^ values for *k*_catCO2_, *k*_catO2_, and *S*_C/O_ (Chen and Spreitzer, 1992; Tcherkez *et al*., 2006; McNevin *et al*., 2007; Tcherkez, 2013). Because *k*_catCO2_ and *k*_catO2_ are first order rate constants they have been represented as

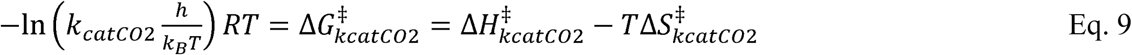

and

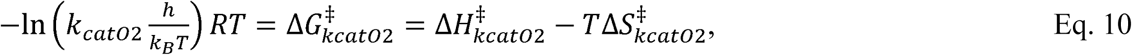

because *S*_C/O_ is the ratio of two first order rate constants (Equation 6) it has been represented as

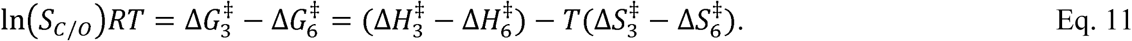

The Δ*G*^‡^ terms in Equations 9, 10, and 11 can be calculated directly from measured values, and the Δ*H*^‡^ and Δ*S*^‡^ terms would describe a linear fit to the temperature response, assuming Δ*H*^‡^ and Δ*S*^‡^ are constant within the temperature range. However, the use of Equations 9, 10, and 11 do not provide information regarding an elementary rate constant or a corresponding energy barrier. Further modeling to estimate individual rate constants from the measured data is described below.

#### Modeling of Radiolabel Data

Each of the rate constants (*k*) in Figure 1 has a corresponding energy of activation (Δ*G*^‡^ from Equation 7), which has a corresponding enthalpic and entropic component (Δ*H*^‡^ and Δ*S*^‡^ from Equation 8). We assumed that the values of Δ*H*^‡^ and Δ*S*^‡^ are constant within the temperature range. Therefore, we fit Michaelis-Menten parameters calculated from elementary rate constants using Equations 2 through 6 to the measured Michaelis-Menten parameters by varying the corresponding Δ*H*^‡^ and Δ*S*^‡^ values. All modeling used the solver function in Excel (2016, Microsoft, Redmon, WA, USA) to minimize the sum of the differences squared between modeled and measured parameters.

The rate constants *k*_8_ (cleavage of carboxylated intermediate) and *k*_9_ (enolization of RuBP) were calculated from measured *k*_catCO2_ values following the calculations of Tcherkez *et al*. (2013) such that *k*_8_/*k*_9_ is 0.83 at 25 °C. The rate constant *k*_10_ (de-enolization) was modeled *assuming k*_9_/*k*_10_ is 0.43 at 25 °C following the calculations of Tcherkez *et al*. (2013), we further assumed that this ratio remained constant with temperature. This allowed for calculation of the rate of *k*_6_ (CO_2_ addition) as the only remaining unknown when fitting measured values of *K*_C_ with Equation 4 assuming *k*_7_ (de-carboxylation) was negligible. After calculating *k*_6_, *k*_3_ (O_2_ addition) was modeled from measured *S*_C/O_ values and Equation 6, assuming rate constants *k*_7_ (decarboxylation) and *k*_4_ (deoxygenation) are negligible. Finally, the rate constant *k*_5_ (cleavage of the oxygenated intermediate) was calculated from measured *K*_O_ values and Equation 7, again assuming *k*_4_ (deoxygenation) was negligible. This process allowed for estimation of the temperature response for k and Δ*G*^‡^ values for each step of the reaction mechanism listed in Equations 2 through 6, with the exception of the decarboxyalation and deoxygenation steps that were assumed negligible (Tcherkez *et al*., 2013; Tcherkez, 2013; Tcherkez, 2016).

#### Modeling of MIMS Data

For the MIMS data, where measurements of *k*_catO2_ were available and non-linearity of Arrhenious plots were observed, the rate constants and corresponding Δ*G*^‡^, Δ*H*^‡^, and Δ*S*^‡^ values were determined differently. The Δ*H*^‡^ and Δ*S*^‡^ values corresponding to the rate constants for *k*_8_ (cleavage of carboxylated intermediate), *k*_5_ (cleavage of oxygenated intermediate), and *k*_9_ (RuBP enolization) were determined by fitting to measured *k*_catCO2_ and *k*_catO2_ values, assuming *k*_8_/*k*_9_ was 0.83 at 25 °C, and using Equations 2 and 3. Because *k*_catCO2_ showed a breakpoint, it is possible that *k*_8_ and *k*_9_ have different temperature responses, with a crossover at approximatly 25 °C. However, *k*_catO2_ also showed a breakpoint at 25 °C and the carbxylated intermediate cleavage rate (*k*_8_) is much greater than the oxygenated cleavage rate (*k*_5_) because measured *k*_catCO2_ values are greater than measured *k*_catO2_. Therefore, a crossover of *k*_5_, *k*_8_, and *k*_9_ at a single temperature is not possible and a breakpoint in *k*_catCO2_ and *k*_catO2_ co-occuring at a single temperature cannot be modeled as a changing rate limiting step. Therefore, we modeled the breakpoint in *k*_catO2_ by allowing *k*_5_ to have separate Δ*H*^‡^ and Δ*S*^‡^ values above and below the breakpoint, and assuming *k*_9_ had the same values of Δ*H*^‡^ and Δ*S*^‡^ above and below the breakpoint. Because *k*_9_ (rate constant of RuBP enolization) temperature response was assumed constant for models of *k*_catO2_, it was also assumed constant when modeling *k*_catCO2_.Therefor, *k*_8_ was allowed to have separate values of Δ*H*^‡^ and Δ*S*^‡^ above and below the breakpoint. The *k*_10_ (rate constant of de-eneolization) was subsequently calculated assuming the ratio *k*_9_/*k*_10_ was 0.43 and constant with temperature. The value *k*_6_ (rate constant of CO_2_ addition) was then calculated from measured *K*_C_ and the approximation of Equation 4 assuming decarboxylation is negligible. This was also done for *k*_3_ (rate constant for O_2_ addition) using *K*_O_ and the approximation of Equation 5 assuming de-oxygenation (*k*_4_) was negligable. It was required that *k*_6_ and *k*_3_ have seperate Δ*H*^‡^ and Δ*S*^‡^ values above and below the breakpoint to model linear Arrhenious plots of *K*_C_ and *K*_O_. This process provided estimates of the temperature response for *k* and Δ*G*^‡^ values for each step of the reaction mechanisms making up the measured Michaelis-Menten parameters (Equations 2 through 6), with the exception of the decarboxyalation and deoxygenation steps, which were assumed negligable.

## RESULTS

### Breakpoints

The Davies test indicated significant breakpoints for the *k*_catCO2_, *k*_catO2_, and *S*_C/O_ temperature response for the MIMS data as well as for the Radiolabel single point measurement of *k*_catCO2_ (Table 1, Figures 2 and 4). Both the Davies test and the maximum likelihood segmented analysis indicated that the breakpoints in these parameters were near 25 **°**C (Table 1). All other parameters showed no breakpoints in their temperature responses for either the MIMS or Radiolabel data sets (Table 1, Figures 2, 3 and 4).

**Figure 2.**
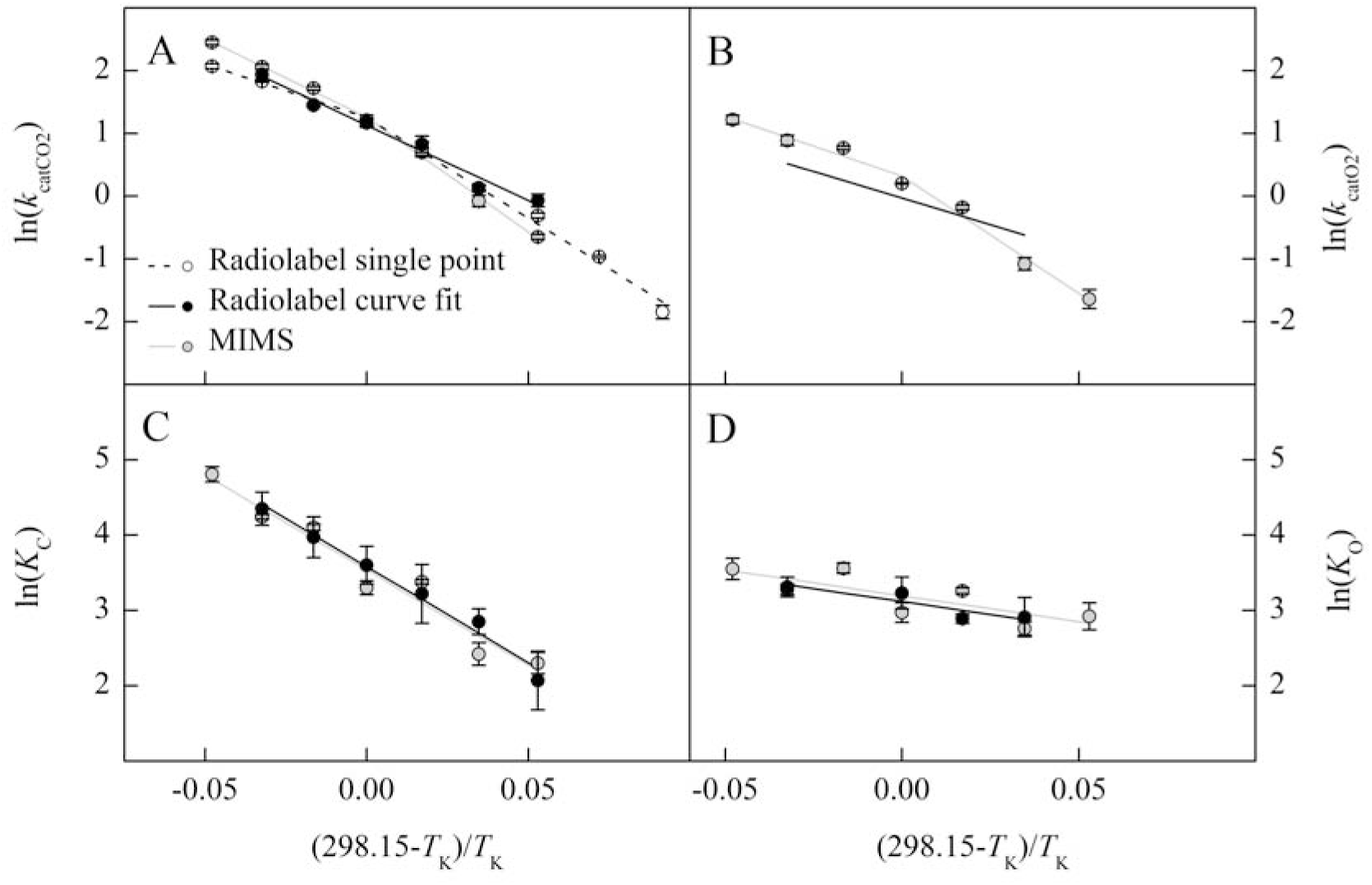
The natural log of Rubisco parameters from *Arabidopsis thaliana* measured using Radiolabel (single point, open circle; curve fit, black circle) and MIMS (grey circle) methods are plotted against the inverse of the temperature in Kelvin offset to a y-intercept of 25 °C. The temperature response of catalytic turnover for CO_2_ (*k*_catCO2_, s^−1^, Panel A), and O_2_ (*k*_catO2_, s^−1^, Panel B), the Michaelis-Menten constant for CO_2_ (*K*C, Pa, Panel C), and O_2_ (*K*_O_, kPa, Panel D) are shown. The lines represent the model fit to the measured data. The Radiolabel *k*_catO2_ model in panel B was determined from the relationship with *S*_C/O_ described by Equation S6.

**Figure 3.**
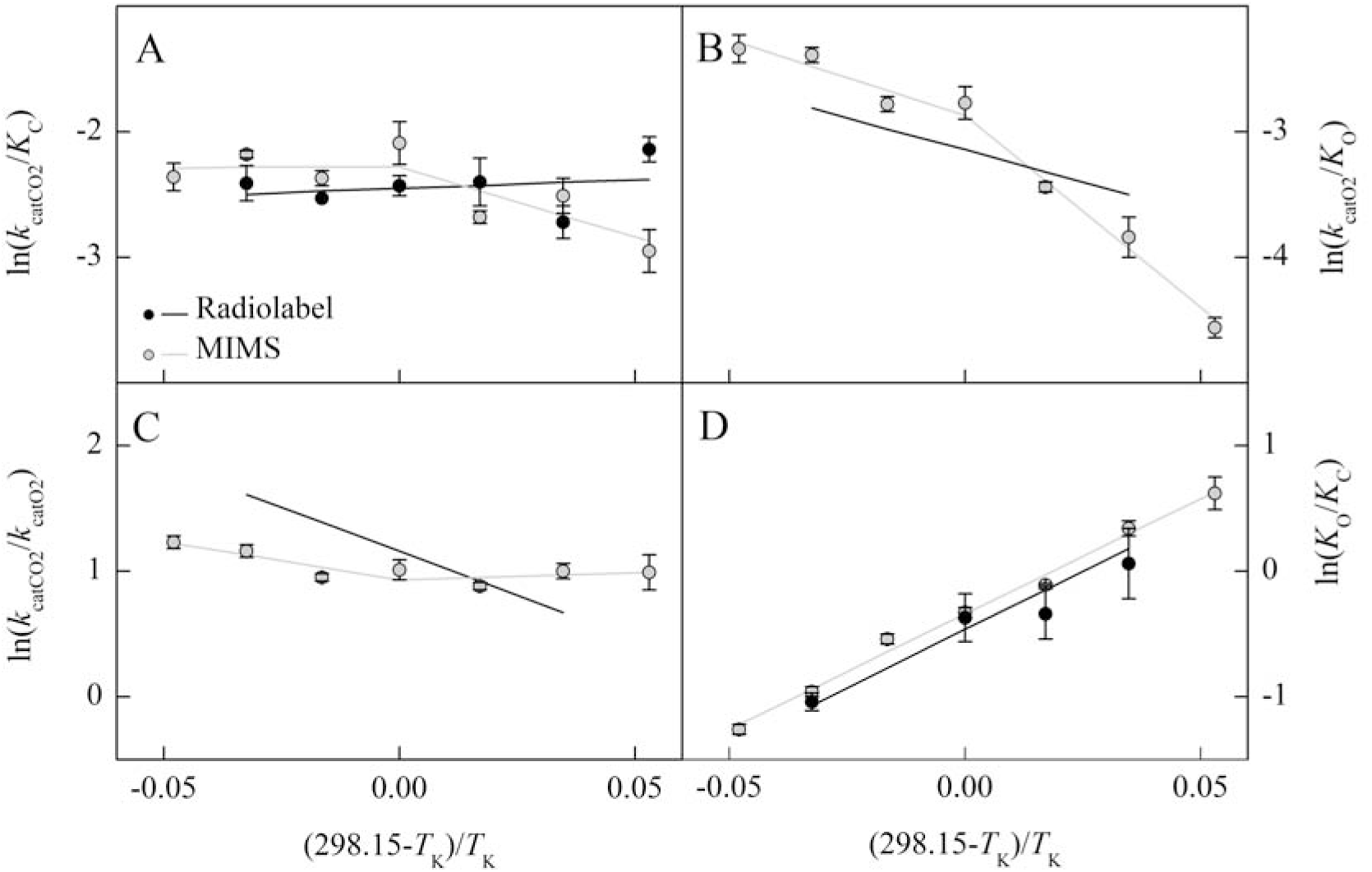
The natural log of the Rubisco parameter ratios from *Arabidopsis thaliana* measured using Radiolabel (black circle) and MIMS (grey circle) are plotted against the inverse of the temperature in Kelvin offset to a y-intercept of 25 °C. The temperature response of the catalytic efficiency of the carboxylation (*k*_catCO2_/*K*C, Panel A) and oxygenation (*k*_catO2_/*K*_O_, Panel B) reactions, catalytic turnover ratio for CO_2_ over O_2_ (*k*_catCO2_/*k*_catO2_, Panel C), and the Michaelis-Menten constant ratio for O_2_ over CO_2_ (*K*_O_/*K*C, Panel D) are shown. Lines represent the combination of models represented in Figure 2 and are not the result of linear regressions to the ratios.

**Figure 4.**
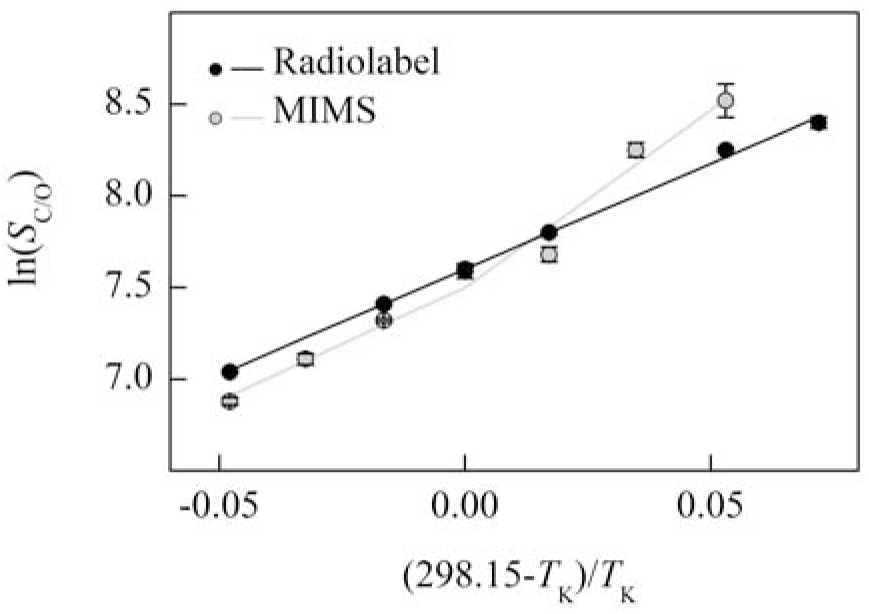
The natural log of Rubisco specificity for CO_2_ over O_2_ (*S*_C/O_) from *Arabidopsis thaliana* measured using Radiolabel (black circle) and MIMS (grey circle) methods are plotted against the inverse of the temperature in Kelvin offset to a y-intercept of 25 °C. The black line represents the model fit to the measured Radiolabel values. The grey line was determined from the relationship of *S*_C/O_ to the parameters presented in Figure 2, described by Equation S6.

**Table 1.** Testing for thermal breaks for all kinetic parameters. Arrhenius plots were examined using the package ‘segmented’ in R (R Core Team, 2013, http://www.R-project.org/), which determines significance of breakpoints in linear models and estimates breakpoint locations as described by Davies (1987). Additionally, breakpoint locations and confidence intervals (CI, lower and upper) were independently estimated using a maximum likelihood test (Muggeo 2003; Muggeo 2008). Where * indicates a p-value < 0.05 for the Davies test and ns = not significant.

| Method | Parameter | Davies Test |  | *Maximum likelihood* |  |  |
| --- | --- | --- | --- | --- | --- | --- |
|  |  | Estimated Breakpoint (°C) | p-value | Estimated Breakpoint (°C) | *CI* (lower) | CI (upper) |
| Radiolabel |  |  |  |  |  |  |
| | $k_{\text{catCO2}}$ single point | 26.8 | * | 25.1 | 5.3 | 36.9 |
| | $k_{\text{catCO2}}$ curve fit | - | ns | | | |
| | $k_{\text{catO2}}$ | - | - | | | |
| | $K_{\text{C}}$ | - | ns | | | |
| | $K_{\text{O}}$ | - | ns | | | |
| | $S_{\text{C/O}}$ | - | ns | | | |
| MIMS |  |  |  |  |  |  |
| | $k_{\text{catCO2}}$ | 25.3 | * | 25.3 | 23.1 | 31.5 |
| | $k_{\text{catO2}}$ | 25.3 | * | 25.5 | 24.3 | 32.6 |
| | $K_{\text{C}}$ | - | ns | | | |
| | $K_{\text{O}}$ | - | ns | | | |
| | $S_{\text{C/O}}$ | 25.4 | * | 25.2 | 15.0 | 27.6 |

### Arrhenius Activation Energies and Modeled Value at 25 °C

The *E*_a_, and *k*_25_ for *k*_catCO2_, *k*_catO2_ (Table 2), *K*_C_, *K*_O_, *S*_C/O_ (Table 3) and ratios of interest (Table 4) were calculated from the linear regressions shown in Figures 2 through 4. For the MIMS derived parameters with breakpoints (*k*_catCO2_, *k*_catO2_, *S*_C/O_), and the Radiolabel single point estimate of *k*_catCO2_ the lower temperatures *E*_a_ values were larger than *E*_a_ values estimated at higher temperatures (Table 2 and 3). Above 25 °C, the *E*_a_ values were similar for all parameters between the Radiolabel and MIMS curve fitting methods. The Radiolabel *E*_a_ for *k*_catCO2_ determined by curve fitting across all temperatures was intermediate to the two *E*_a_ values estimated above and below the breakpoint from the single point Radiolabel data. The *k*_25_ values for *k*_catCO2_ estimated from Radiolabel and MIMS methods were not different from each other, but were larger than the *k*_25_ for *k*_catO2_ determined by MIMS (Table 2). The *E*_a_ and *k*_25_ values for *K*_C_ and *K*_O_ were not significantly different between methods (Table 3). However, the MIMS *S*_C/O_ measured from 10 to 25 °C had a lower (more negative) *E*_a_ value than the MIMS *S*_C/O_ *E*_a_ value measured from 25 to 40 °C and the Radiolabel *S*_C/O_ *E*_a_ value (Table 3).

**Table 2.** Comparison of *k*_25_ and *E*_a_ values for *k*c_a_t measurements from the different methods. The *k*_25_ and *E*_a_ values are the mean of three to four replicates, calculated from linear regressions of Arrhenius plots. The temperature ranges for each regression were determined by segment analysis. Superscripts indicate significant differences between groups (Tukey HSD, p < 0.05).

| Method | Temperature (°C) | Parameter | $k_{25}$ | $E_a$ |
| --- | --- | --- | --- | --- |
| Radiolabel |  |  |  |  |
| single point | 0 - 25 | $k_{catCO_2}$ (s<sup>-1</sup>) | 3.50 ±0.20<sup>A</sup> | 79.53 ±2.03<sup>a</sup> |
|  | 25 - 40 |  | -- -- | 42.11 ±3.45<sup>c</sup> |
| curve fit | 10 - 35 |  | 3.10 ±0.07<sup>A</sup> | 59.64 ±3.93<sup>b</sup> |
| MIMS | 10 - 25 |  | 3.53 ±0.25<sup>A</sup> | 90.36 ±1.03<sup>a</sup> |
|  | 25 - 40 |  | -- -- | 62.20 ±2.68<sup>b</sup> |
| | 10 - 25 | $k_{catO_2}$ (s<sup>-1</sup>) | 1.38 ±0.05<sup>B</sup> | 92.95 ±7.31<sup>a</sup> |
|  | 25 - 40 |  | -- -- | 47.11 ±2.33<sup>bc</sup> |

**Table 3.** Comparison of *K*_C_, *K*_O_, *S*_C/O_ parameters *k*_25_ and *E*_a_ resulting from the different methods are shown. No differences were observed in *k*_25_ between methods. No differences were observed in *E*_a_ values for *K*_C_ and *K*_O_ values between methods (ANOVA). The superscripts next to the *E*_a_ values indicate significant differences for the *S*_C/O_ values (Tukey HSD, p < 0.05).

| Method | Temperature Range (°C) | Parameter | $k_{25}$ (Pa) | | $E_a$ (kJ mol<sup>-1</sup>) | |
| --- | --- | --- | --- | --- | --- | --- |
| Radiolabel | 10 - 35 | $K_C$ | 36 | ±2 | 63.09 | ±6.23 |
| MIMS | 10 - 40 |  | 34 | ±1 | 62.62 | ±3.44 |
| Radiolabel | 15 - 35 | $K_O$ | 23100 | ±3430 | 16.89 | ±2.59 |
| MIMS | 10 - 40 |  | 24400 | ±701 | 17.01 | ±2.48 |
| Radiolabel | 5 - 40 | $S_{C/O}$ | 2003 | ±22 | -28.66 | ±0.51<sup>b</sup> |
| MIMS | 10 - 25 |  | 1814 | ±117 | -48.19 | ±4.17<sup>a</sup> |
|  | 25 - 40 |  | -- | -- | -30.51 | ±6.41<sup>b</sup> |

The *E*_a_ value for the carboxylation efficiency (*k*_catCO2_/*K*_C_) below 25 °C was significantly different from zero for the MIMS method, where the carboxylation efficiency increased with temperature; however, above 25 **°**C the *E*_a_ was not significantly different from zero (Table 4). The MIMS *E*_a_ for oxygenation efficiency (*k*_catO2_/*K*_O_) was significantly different from zero above and below 25 °C (Table 4). The *E*_a_ for the ratio of catalytic rates (*k*_catCO2_/*k*_catO2_) measured by MIMS was only significantly different than zero above 25 °C (Table 4). The *E*_a_ for *K*_O_/*K*_C_ was significantly different from zero for both Radiolabel and MIMS methods (Table 4).

**Table 4.**
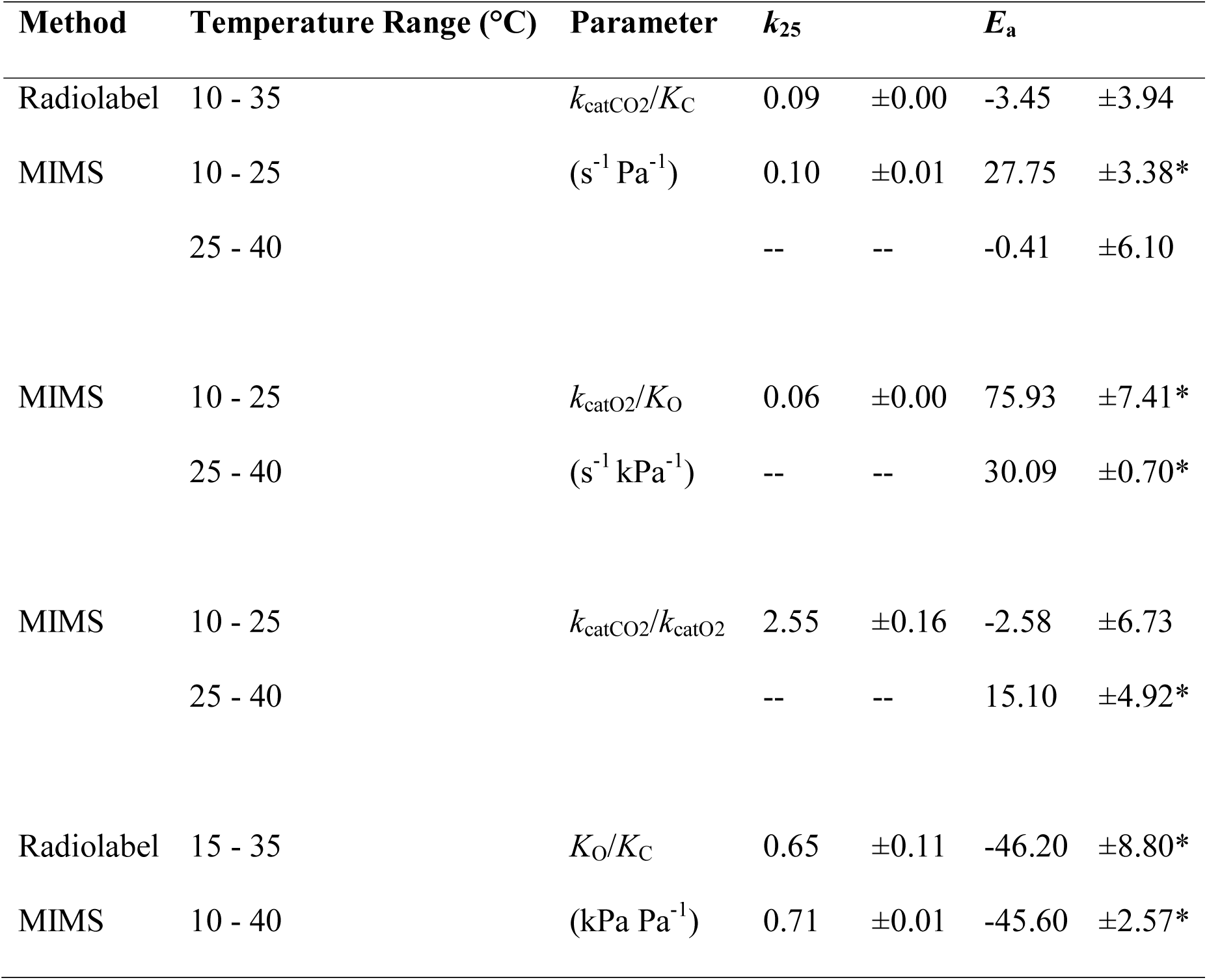
The *E*_a_ and *k*_25_ parameters for *k*_catCO2_/*K*C, *k*_catO2_/*K*O, *k*_catCO2_/*k*_catO2_, and *K*O/*K*C ratios. The *E*_a_ parameters were tested to determine if they were significantly different than zero (t-test), where the * next to the *E*_a_ values indicates a p-value below 0.05.

| Method | Temperature Range (°C) | Parameter | $k_{25}$ | | $E_a$ | |
| --- | --- | --- | --- | --- | --- | --- |
| Radiolabel | 10 - 35 | $k_{\text{catCO}_2}/K_C$ | 0.09 | ±0.00 | -3.45 | ±3.94 |
| MIMS | 10 - 25 | (s<sup>-1</sup> Pa<sup>-1</sup>) | 0.10 | ±0.01 | 27.75 | ±3.38* |
|  | 25 - 40 |  | -- | -- | -0.41 | ±6.10 |
| MIMS | 10 - 25 | $k_{\text{catO}_2}/K_O$ | 0.06 | ±0.00 | 75.93 | ±7.41* |
|  | 25 - 40 | (s<sup>-1</sup> kPa<sup>-1</sup>) | -- | -- | 30.09 | ±0.70* |
| MIMS | 10 - 25 | $k_{\text{catCO}_2}/k_{\text{catO}_2}$ | 2.55 | ±0.16 | -2.58 | ±6.73 |
|  | 25 - 40 |  | -- | -- | 15.10 | ±4.92* |
| Radiolabel | 15 - 35 | $K_O/K_C$ | 0.65 | ±0.11 | -46.20 | ±8.80* |
| MIMS | 10 - 40 | (kPa Pa<sup>-1</sup>) | 0.71 | ±0.01 | -45.60 | ±2.57* |

### Modeling *k* and Δ*G*^‡^

Above 25 °C the Δ*G*3^‡^-Δ*G*6^‡^ for *S*_C/O_ from Radiolabel and MIMS (Fig. 5) are similar to previous calculations for C_3_ species reported by Tcherkez *et al*. (2006). However, the MIMS entropy difference between O_2_ and CO_2_ addition (Δ*S*3^‡^-Δ*S*6^‡^, slope of line in Fig. 5, see Eq. 11, Supp. Table 1), from data colleted below 25 °C appear more similar to the Δ*S*3^‡^-Δ*S*6^‡^ of red algae rather than higher plants, when compared to data presented in Tcherkez *et al*. (2006).

**Figure 5.**
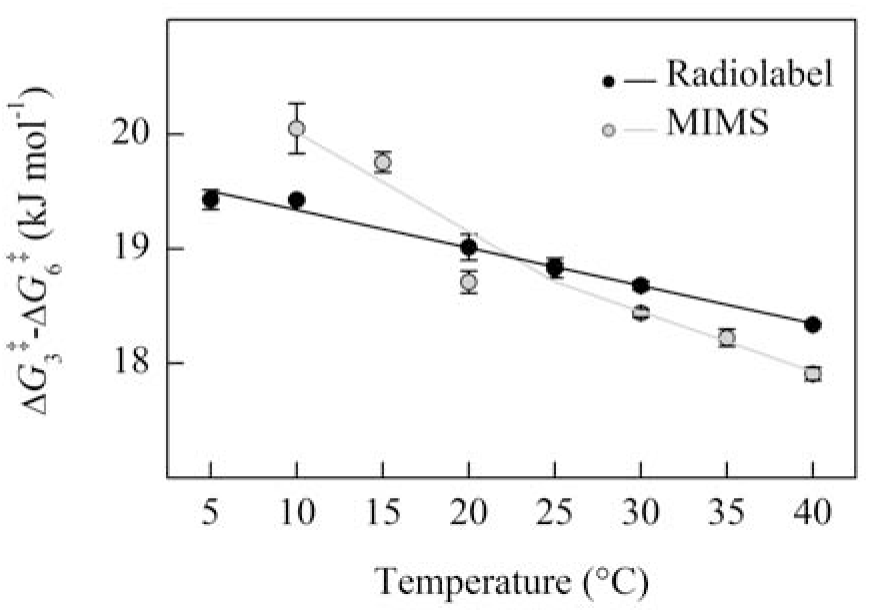
The temperature response of Δ*G*3^‡^-Δ*G*6^‡^ calculated from the data presented in Figure 4. Both measurement methods show a decrease with temperature. Solid black circles are the mean of four replicates measured using Radiolabel, filled grey circles are the means from three replicates using MIMS, standard error is shown. The solid lines indicate the linear regression fit to calculated values.

The free energy of activation associated with *k*_catCO2_ (Δ*Gk*_catCO2_ ^‡^) plotted against temperature, increased linearly for the Radiolabel curve fit method, while the Δ*Gk*_catCO2_ ^‡^ calculated from MIMS measurements decreased from 10 to 25 °C and then increased from 25 to 40 °C (Fig. 6). A similar temperature response was also observed for MIMS Δ*Gk*_catO2_ ^‡^, although the absolute values of Δ*Gk*_catO2_ ^‡^ are larger than Δ*Gk*_catCO2_ ^‡^ as evident by a lower *k*_catO2_ compared to *k*_catCO2_ at all temperatures (i.e. larger energy barriers result in slower reactions). The slope of Δ*Gk*_catCO2_ ^‡^ values presented in Figure 6 (equivalent to the entropy term Δ*S*_kcatCO2_ ^‡^, Supp. Table 2) calculated for Radiolabel and MIMS above 25 °C are slightly larger than those reported for *Nicotiana tabacum* (McNevin *et al*., 2007). The MIMS Δ*S*_kcatCO2_ ^‡^ and Δ*Sk*_catO2_ ^‡^ showed a sign change above and below the breakpoint (negative slope to positive slope, Fig. 6, Supp. Table 2).

**Figure 6.**
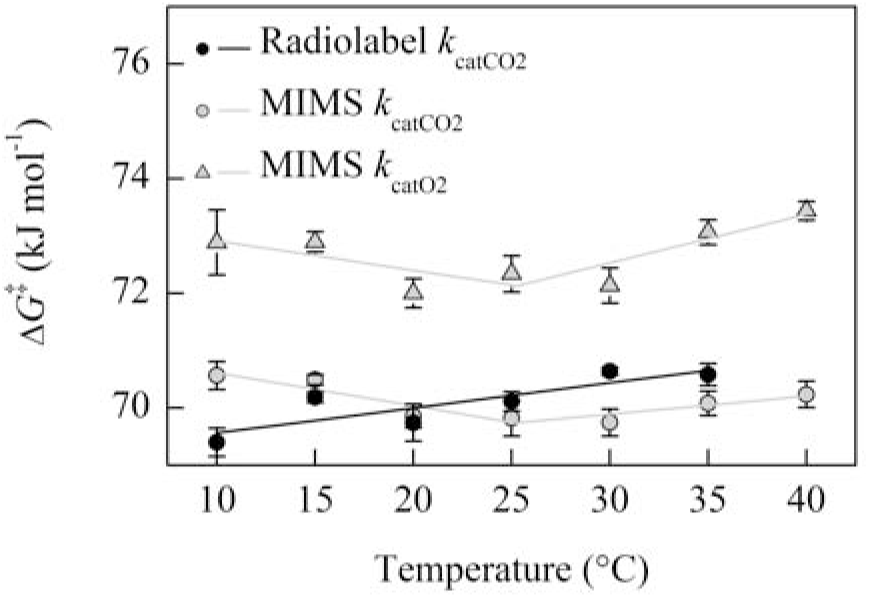
The temperature response of Δ*Gk*_catCO2_ ^‡^ for MIMS and Radiolabel methods, and Δ*Gk*_catO2_ ^‡^ for MIMS calculated from the data presented in Figure 2. Two regressions were fit to the MIMS data on either side of the 25 °C breakpoint, a single regression is fit to the Radiolabel data. Solid black circles are the mean of three replicates measured using Radiolabel, filled grey circles are the means from three replicates using MIMS, standard error is shown.

Temperature responses of the rate constants (*k*) and corresponding energy barriers of the transition states (Δ*G*^‡^) are shown in Figure 7, while the modeled Δ*H*^‡^ and Δ*S*^‡^ values are presented in Suppemental Table 3. Calculations of elementary rate constants and corresponding Δ*G*^‡^ are similar to previous calculations at 25 °C from Tcherkez (2013) and Tcherkez (2016). In order to model breakpoints in the MIMS *k*_catCO2_, *k*_catO2_, and *S*_C/O_ parameters, breakpoints are neeeded in the rate constants for the cleavage (*k*_8_ and *k*_5_) and for gas addition (*k*_6_ and *k*_3_). This is required because it was not possible to model a simultaneous change in the rate limiting step for both the *k*_catCO2_ and *k*_catO2_ parameter (Supp. Fig. 2). This further required that breakpoints were needed in the rate constants for CO_2_ and O_2_ addition (*k*_6_ and *k*_3_, respectively) to maintain the observed linearity for *K*_C_ and *K*_O_ Arrhenius plots (Fig. 2).

**Figure 7.**
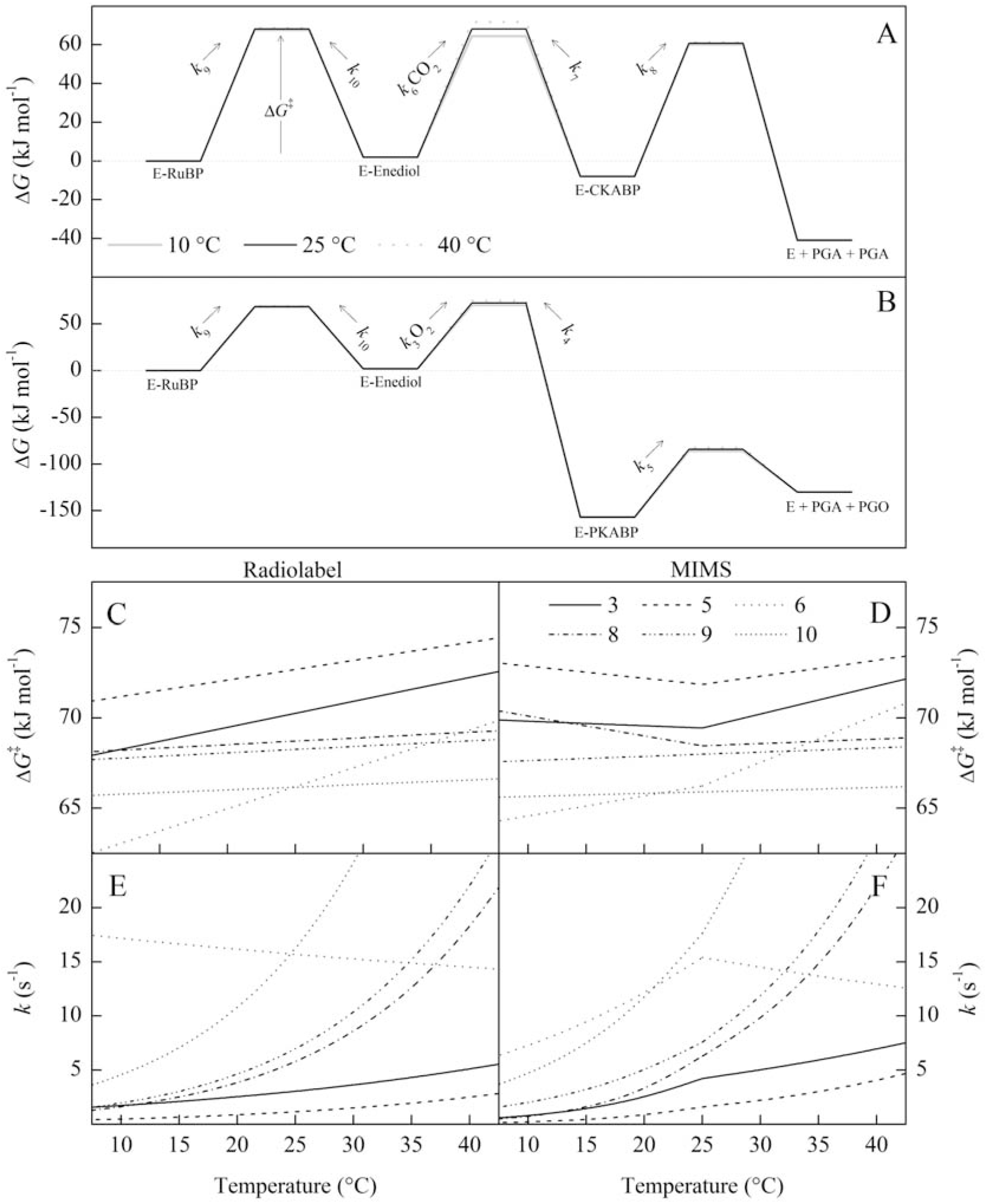
A kinetic energy barrier diagram showing the modeled temperature responses of the energy barrier to the transition state (Δ*G*^‡^) and the corresponding first order rate constant *k*. The Δ*G*^‡^ and *k* are indicated by the numbered step of the reaction following Figure 1. The assumptions made for this model are stated in the methods section. For steps 3 and 6 (O_2_ and CO_2_ addition, respectively), the rate constants were multiplied by ambient concentrations O_2_ (21 kPa) and CO_2_ (41 Pa) as a pseudo-first order approximation for comparison to the other rate constants and to calculate their respective Δ*G*^‡^. For the bottom figure, the left-hand column is modeled on the Radiolabel data and the right-hand column on the MIMS data so that comparisons between continuous and breakpoint temperature responses can be made. The values for intermediates were taken from Tcherkez (2013) for Panel A and Tcherkez (2016) for Panel B and assumed to remain constant with temperature.

## DISCUSSION

### Temperature Responses of Rubisco Michaelis-Menten Kinetic Parameters

The Rubisco kinetic parameters for *Arabidopsis thaliana* measured with the Radiolabel and MIMS curve fitting methods were similar at and above 25 °C. Additionally, the modeled 25 °C values (*k*_25_) and Arrhenius activation energy (*E*_a_) above 25 °C agree with many of the literature values for other C_3_-type Rubiscos, including *in vitro* and *in vivo* measurements of *A. thaliana* (Flexas *et al*., 2007; Whitney *et al*., 2011; Walker *et al*., 2013; Weise *et al*., 2015; Galmés *et al*., 2016). Although, previous reports of Rubisco specificities for CO_2_ over O_2_ (*S*_C/O_) at 25 °C vary widely for C_3_ species, including for *A. thaliana* which range from below 2125 to above 2655 (Pa Pa^−1^; Flexas *et al*., 2007; Whitney *et al*., 2011; Walker *et al*., 2013; Weise *et al*., 2015).

Galmés *et al*. (2016) highlighted contradictory trends in the temperature response of *K*_O_ when measured by *in vitro* assay; either increasing or decreasing with temperature (when expressed in units of molarity; converting between molarity and partial pressure changes the temperature response because the solubility of O_2_ decreases with temperature). Here, both the Radiolabel and MIMS method show increases in *K*_O_ with temperature, with lower *E*_a_ values compared to *K*_C_. The two data sets presented here confirm trends from the growing literature on C_3_ Rubisco temperature responses, at least for values measured above 25 °C.

Alternatively, below 25 °C the Radiolabel and MIMS derived parameters had different temperature responses where the Arrhenius plots of MIMS determined *k*_catCO2_, *k*_catO2_, and *S*_C/O_ were non-linear (Fig. 2, Fig. 4). Different temperature responses at high and low temperatures were interpreted as breakpoints for these kinetic parameters at 25 °C (Fig. 2). However, for the Radiolabel curve fit data all kinetic parameters appeared sufficiently linear. This could suggest methodology artifacts; however, it is difficult to identify methodological errors that may give rise to breakpoints given that they have also been observed by different laboratories using varying methods and species (Badger and Collatz, 1977; Sage 2002, Kubien *et al*., 2003; Sharwood *et al*., 2016).

### Evidence for Breakpoints in the Literature

Björkman and Pearcy (1970) first identified a thermal breakpoint occurring in the temperature response of *V*cm_a_x from two Atriplex species. However, in the same publication they determined that the apparent breakpoints were caused by non-saturating or inhibitory bicarbonate concentrations at varying temperatures and, when corrected, they obtained sufficiently linear Arrhenius plots. Subsequently, Badger and Collatz (1977) identified breakpoints in *k*_catCO2_, *k*_catO2_, and *K*_C_ occuring at 15 °C, with sufficiently linear Arrhenius plots of *K*_O_. While Badger and Collatz (1977) did not discuss *S*_C/O_, using Equation S6 with their data suggests a breakpoint in *S*_C/O_. Badger and Colloatz (1977) hypothesized that breakpoints were the result of possible changes in enzyme conformation which change the rate limiting step of the reaction mechanism. Sage (2002) idenified breakpoints in *k*_catCO2_ at 22 °C for *Oryza sativa*, but observed linear Arrhenius plots for other species. Furthermore, Kubien *et al*. (2003) also observed a breakpoint in *k*_catCO2_ between 12 and 18 °C for *Flaveria bidentis*, and suggested it was caused by deactivation of the enzyme at low temperature, possibly by dissociation of the haloenzyme. These analyses generally identified *E*_a_ values above the breakpoint similar to what is often reported in the literature for the temperature response of Rubisco (*E*_a_ for *k*_catCO2_ ~60 kJ mol^−1^), with larger *E*_a_ values at low temperatures.

Recently, Sharwood *et al*. (2016) suggested breakpoints occuring at 25 °C for *k*_catCO2_ in eleven Panicoid grasses and tobacco. While *E*_a_ values at lower temperatures remain larger than the *E*_a_ values at higher temperatures, Sharwood *et al*.’s (2016) findings differs from the previous breakpoint publications, because the *E*_a_ below the breakpoint is around the expected value (~60 kJ mol^−1^) and the *E*_a_ above the breakpoint is lower than expected (~30 kJ mol^−1^). Sharwood *et al*. (2016) did not observe breakpoints in *S*_C/O_, but it is worth noting that they calculated *S*_C/O_ from a separate assay from *k*_catCO2_ using the ratio of ^3^H-glycerate to ^3^H-glycolate as described here for the Radiolabel method. They also did not measure the temperature responses of *K*_C_, *K*_O_ or *k*_catO2_ limiting direct comparisons to the data presented here.

### Radiolabel Single Point *k*_catCO2_ Breakpoint

The Radiolabel single point method reported here utilized a single bicarbonate concentration with temperature (11 mM). This method resulted in a breakpoint, having an *E*_a_ value of 79.5 kJ mol^−1^ at low temperatures, and 42.1 kJ mol^−1^ at higher temperatures (Fig. 2, Table 2). Similar to Björkman and Pearcy (1970), the linear Arrhenius plot (Radiolabel curve fit) has an *E*_a_ value intermediate to the two *E*_a_ values determined when using a single bicarbonate concentration (~59.6 kJ mol^−1^, Fig. 2, Table 2). Because Björkman and Pearcy (1970) suggested that there could be inhibition at low temperature and sub-saturating concentrations at high temperature, we plotted the predicted CO_2_ concentration achieved by 11 mM NaHCO_3_ at each temperature against the measured and modeled CO_2_ response of the enzyme determined by both Radiolabel and MIMS curve fitting methods (Supp. Fig. 1). The CO_2_ concentration provided by the 11 mM NaHCO_3_ is less saturating at higher temperatures because the *K*_C_ of Rubisco increases with temperature and the pK_a_ temperature response favors HCO_3_^−^ at higher temperatures (Supp. Fig. 1).

From the data presented here, the CO_2_ concentration appears saturating at 10 and 15 °C, but becomes increasingly less saturating at higher temperatures, as indicated where the shaded area intersects the modeled CO_2_ response. This suggest the lower *E*_a_ value of the single point method at high temperatures could be caused by sub-saturating CO_2_ concentrations. The sub-saturating CO_2_ concentrations is likely due to both an increase in *K*_C_ with temperature and the predicted concentration of CO_2_ decreases with temperature given the temperature response of the pK_a_ (Harned and Bonner, 1945). Alternatively, an inhibitory concentrations of CO_2_ was not observed under any of the measurement conditions.

### MIMS *k*_catCO2_, *k*_catO2_, and *S*_C/O_ Breakpoints

The non-linearity of Arrhenius plots of *k*_catCO2_, *k*_catO2_, and *S*_C/O_ for the MIMS data were interpreted as 25 °C breakpoints. Badger and Collatz (1977) also observed breakpoints in *k*_catCO2_, *k*_catO2_, and *S*_C/O_; however, they observed an additional thermal breakpoint in *K*_C_, which was not observed with the MIMS data presented here. As *S*_C/O_ is a ratio of *k*_catCO2_, *K*_C_, *K*_O_, and *k*_catO2_ (Eq. S6), the differences in *S*_C/O_ breakpoints between Badger and Collatz (1977) and our MIMS data could suggest different mechanisms driving the thermal response of *S*_C/O_. Furthermore, no breakpoint in *S*_C/O_ has been observed in any study using the ^3^H-RuBP method.

The breakpoints observed in MIMS *k*_catCO2_ and *k*_catO2_ are unlikely to be caused by insufficient or inhibitory CO_2_ concentrations, as is possible for the breakpoint observed in the Radiolabel single point *k*_catCO2_ measurement, as sub-saturation or inhibition should be evident in the CO_2_ response curves (Supp. Fig. 1). A breakpoint in both *k*_catCO2_ and *k*_catO2_ could be caused by deactivation of the enzyme as was suggested by Kubien *et al*. (2003). However, deactivation is unlikely to change the *k*_catCO2_/*k*_catO2_ temperature response as was observed in Figure 3C, because both catalytic rates are expected to be affected in the same way by deactivation. Alternatively, the observed breakpoints in MIMS could be related to methodology as the Radiolabel Arrhenius plots presented here for *k*_catCO2_ and *S*_C/O_ were sufficiently linear.

Nevertheless, breakpoints have persisted in the Rubisco literature for over forty years without sufficient explanation and warrant further investigations into their underlying causes. Badger and Collatz (1977) suggested changes in the rate-limiting step of the reaction mechanism brought about by conformational changes. If the elementary rate constants defining a specific parameter have different temperature responses then this could cause breakpoints if they crossover causing a change in rate limiting step. The discussion below utilizes the currently accepted reaction mechanism of Rubisco (Fig. 1) and transition state theory to explore breakpoints as a function of changes in energy barriers to elementary reactions.

### Rubisco Reaction Mechanisms and Breakpoints

#### Radiolabel modelling

For the Radiolabel data, where all Arrhenius plots were sufficiently linear, a model of how the energy barriers for the Rubisco reaction mechanism change with temperature is presented in Figure 7, Panel C and E, and depicted as a kinetic energy barrier diagram in Panel A and B. As previously suggested by Tcherkez (2013) the *k*_catCO2_ and *k*_catO2_ values can be modeled assuming identical temperature responses for the rate of enolization (*k*_9_), and cleavage for the carboxylated intermediate (*k*_8_) and oxygenated-intermediate (*k*_5_). Interestingly, the modeled addition of CO_2_ (Δ*G*6^‡^) had high entropic cost leading to a decreasing temperature response for the rate of CO_2_ addition (*k*_6_), suggesting the reaction becomes slower with increasing temperature. Additionally, the increase in the energy barrier for CO_2_ addition (Δ*G*6^‡^) is greater than that for O_2_ addition (Δ*G*3^‡^) such that the ratio *k*_6_/*k*_3_ decreased with temperature. This fits with the observation that Δ*G*3^‡^-Δ*G*6^‡^ decreases with temperature (Fig. 5). While our model for both CO_2_ and O_2_ addition has positive entropy of the transition states, the greater entropic cost for CO_2_ addition could be the cause of *S*_C/O_ decreases with temperature, more than would be assumed if Δ*G*3^‡^-Δ*G*6^‡^ remained constant with temperature.

#### MIMS modeling

For the MIMS data, the breakpoints observed in *k*_catCO2_ and *k*_catO2_ could be due to changes in rate limiting step as suggested by Badger and Collatz (1977). For example, *k*_catCO2_ is a function of the rate of cleavage of the carboxylated-intermediate (*k*_8_) and the rate of RuBP enolization (*k*_9_). This would mean that *k*_8_ and *k*_9_ have different temperature response such that they crossover around the breakpoint observed at 25 °C. However, modeling this change in rate limiting steps due to different temperature responses cannot simultaneously explain the observed breakpoint in *k*_catCO2_ and *k*_catO2_, because the value of *k*_5_ defining the cleavage of the oxygenated intermediate is lower than *k*_8_. This means that *k*_9_ cannot crossover both *k*_8_ and *k*_5_ at 25 °C (Supp. Fig. 2). Therefore, we proposed that rather than a crossover between elementary rate constants, a conformation change in the enzyme could change the temperature response for the cleavage reactions for both carboxylated (*k*_8_) and oxygenated (*k*_5_) intermediates (Fig. 7). The needed change in cleavage reactions (*k*_8_ and *k*_5_) to model a breakpoint suggests a positive entropy for the transition state below the breakpoint (decreasing Δ*G*5^‡^ with temperature) and a negative entropy of the transitions state above the breakpoint (increasing Δ*G*5^‡^ with temperature). While it seems unlikely that such an entropy change could be driven by a conformation change in the enzyme brought about by such minimal changes in temperature, a similar change in entropy for *k*_catCO2_ was observed between wild type *Nicotiana tabacum* and a mutant (L335V) Rubisco (McNevin *et al*., 2007). The amino acid substitution in the mutant was suggested to affect the loop that closes over RuBP as it is bound. This could suggest that the entropy changes to the cleavage reactions (*k*_5_ and *k*_8_) proposed here maybe possible given enzyme conformational changes with temperature.

The MIMS data also indicates a breakpoint in *S*_C/O_ suggesting larger *E*_a_ values at low temperatures compared to higher temperatures, therefor the term Δ*G*3^‡^-Δ*G*6^‡^ was modeled with a non-linear temperature response (Fig. 5). As *S*_C/O_ can be approximated as *k*_6_/*k*_3_, this could suggest a breakpoint in the temperature response of CO_2_ addition (*k*_6_), O_2_ addition (*k*_3_), or both. The individual values for *k*_6_ and *k*_3_ cannot be derived from *S*_C/O_ measurements; however, in order for the observed constant temperature response of *K*_C_ and *K*_O_ to remain constant with temperature the cleavage reactions discussed above need to be offset by breakpoints in both *k*_6_ and *k*_3_ (Fig. 7F). Therefore, to model the reaction mechanism suggested by MIMS measurements, breakpoints in four elementary rate constants are needed to describe the breakpoints in *k*_catCO2_, *k*_catO2_, and *S*_C/O_ but not in *K*_C_ or *K*_O_.

The modeling presented here is largely based on isotope exchange studies, which suggest similar energy barriers between enolization (Δ*G*9^‡^) and cleavage (Δ*G*8^‡^). However, these measurements have been limited to 25 °C (Van Dyk and Schloss, 1986; Tcherkez *et al*., 2013) and extension of isotope exchange studies to temperature responses would help constrain how the elementary rate constants vary with temperature. Contrary to the above proposal that the cleavage transition state (*k*_8_) undergoes changes above and below 25 °C, is that Rubisco discrimination against ^13^CO_2_ is believed to remain constant with temperature (Christeller *et al*., 1976). If the rate of cleavage (*k*_8_) decreases, then the decarboxylation reaction (*k*_7_) may increase, or the ratio *k*_7_/*k*_8_ could increase, which would change Rubisco discrimination against ^13^CO_2_. Furthermore, the above modeling relies on the assumption that decarboxylation (*k*_7_) was negligible at all temperatures; therefore, changes in fractionation with temperature for an enzyme exhibiting breakpoints should help test the validity of these assumptions.

## CONCLUSION

The measured temperature responses of Rubisco kinetic parameters were consistent between methods at and above 25 °C; however, there were thermal breakpoints at ~ 25 °C in the MIMS dataset for *k*_catCO2_, *k*_catO2_, and *S*_C/O_. Additionally, the Radiolabel method using a single bicarbonate concentration showed a breakpoint for *k*_catCO2_ at 25 °C but the curve fitting did not, suggesting this breakpoint was caused in part by non-saturating CO_2_ concentrations at higher temperatures. Previous studies suggest that breakpoints are caused by either a change in the rate limiting step of the reaction mechanism or deactivation of the enzyme at low temperatures. By modeling elementary steps of the reaction mechanism, we showed that a simple change in rate limiting step is not sufficient to explain simultaneous breakpoints in both *k*_catCO2_ and *k*_catO2_, and that breakpoints in the elementary rate constants are likely needed. Additionally, it is unclear how deactivation would cause the observed breakpoint in *S*_C/O_. Moving forward, the temperature response of isotopic substitutions experiments would advance our understanding of how elementary rate constants change in relation to one another with temperature.

## ACKNOWLEDGEMENT

This work was supported by the Division of Chemical Sciences, Geosciences, and Biosciences, Office of Basic Energy Sciences, Department of Energy (grant no. DE–FG02– 09ER16062), the National Science Foundation (Major Research Instrumentation grant no. 0923562), the National Science and Engineering Research Council of Canada (Discovery grant no. 327103; and PGS-D scholarship to APC), and the Seattle chapter of the Achievement Rewards for College Scientists Foundation (R.A.B.). A.B.C. and D.S.K. proposed the original concept and design for the project; R.A.B. and A.P.C. performed the experiments and data analysis; R.A.B. wrote the article with the contributions of all the authors; A.B.C. supervised and complemented the writing. We would like to thank Chuck Cody for maintaining plant growth facilities and current members of Cousins Lab for helpful and insightful discussions.

